# Transfer Elastic Net for Developing Epigenetic Clocks for the Japanese Population

**DOI:** 10.1101/2024.05.19.594899

**Authors:** Yui Tomo, Ryo Nakaki

**Affiliations:** Keio University School of Medicine; Rhelixa Inc.

**Keywords:** Epigenetic clock, DNA methylation, Transfer learning, Elastic Net, Japanese population

## Abstract

**Motivation:** The epigenetic clock evaluates human biological age based on DNA methylation patterns. It takes the form of a regression model where the methylation ratio at CpG sites serves as the predictor, and chronological or adjusted age as the response variable. Due to the large number of CpG sites considered as candidate explanatory variables and their potential correlation, Elastic Net is commonly used to train the regression models. However, existing standard epigenetic clocks, trained on multiracial data, may exhibit biases due to genetic and environmental differences among specific racial groups. The development of epigenetic clocks suitable for a single-race population typically necessitates the collection of hundreds to thousands of samples to measure DNA methylation and other biomarkers, which costs a lot of time and money. Consequently, a method for developing accurate epigenetic clocks with relatively small sample sizes is needed.

**Results:** We propose Transfer Elastic Net, a transfer learning approach that uses the parameter information from a linear regression model trained with the Elastic Net to estimate another model. Using this method, we constructed Horvath’s, Hannum’s, and Levine’s types of epigenetic clocks using DNA methylation data from blood samples of 143 Japanese subjects. The data were transformed through principal component analysis to obtain more reliable clocks. The developed clocks demonstrated the smallest prediction errors compared to both the original clocks and those trained with the Elastic Net on the same Japanese data. Furthermore, the bias relative to the original clocks was reduced. Thus, we successfully developed epigenetic clocks that are well-suited for the Japanese population. Transfer Elastic Net can also be applied to develop epigenetic clocks for other specific populations, and is expected to be applied in various fields.

**Availability:** https://github.com/t-yui/TransferENet-EpigeneticClock

## 1 Introduction

The epigenetic clock calculates the biological age of an individual based on DNA methylation information (Horvath, 2013). DNA methylation, an epigenetic modification, involves the binding of a methyl group to the cytosine base of a DNA molecule. The epigenetic clock specifically focuses on methylation at the guanine-cytosine base pairs (CpG sites; CpGs), which are inherited during cell division. Although DNA methylation does not alter the DNA sequence itself, it regulates gene expression and is influenced by a variety of genetic and environmental factors. The biological age predicted by the epigenetic clock has gained more and more attention in clinical medicine as an important aging indicator. For instance, it can assess the risk of age-related diseases and serve as an endpoint in clinical trials for anti-aging interventions. Various epigenetic clocks have been developed, differing in the tissue source of the biopsy and the statistical methods used to construct the model, such as Hannum’s clock, Zhang’s clock, and Alfonso’s clock (Hannum et al., 2013; Zhang et al., 2019; Alfonso and Gonzalez, 2020). While these clocks are trained to predict chronological age (first-generation clocks), others, like Levine’s PhenoAge and Lu’s GrimAge, incorporate adjusted age considering age-related risk factors for death (second-generation clocks) (Levine et al., 2018; Lu et al., 2019a, Lu et al. 2022).

The epigenetic clock is typically developed as a linear regression model with the measured DNA methylation ratio of CpGs as the explanatory variable and chronological age as the target variable. Given a large number of measured CpGs relative to the sample size, typically ranging from tens to hundreds of thousands, estimating regression coefficient parameters using ordinary least squares or maximum likelihood estimation becomes challenging. Consequently, regularization methods are often used to construct epigenetic clocks. Assuming that only a fraction of the CpGs contribute to age prediction, the Elastic Net, a sparse estimation approach, has become the standard method for developing epigenetic clocks (Zou and Hastie, 2005). Similar to the least absolute shrinkage and selection operator (Lasso), the Elastic Net promotes parameter sparsity by adding a penalty term based on the *ℓ*_1_ norm of the parameters to the log-likelihood function (Tibshirani, 1996). This feature allows CpGs that strongly predict age to be automatically selected and included in the model. The Elastic Net applies an additional *ℓ*_2_-norm penalty, which induces several advantageous properties to the estimator. While Lasso is limited in the number of variables it can select, which is bound by the smaller of the sample size or parameter dimensions, the Elastic Net faces no such limitation. Moreover, the Elastic Net can manage highly correlated explanatory variables through its grouping effect, which states the regression coefficients of strongly correlated variables have small differences (Zhou, 2013). Given that the number of CpGs that significantly contribute to age prediction is typically unknown and may be strongly correlated, it is preferable to use a method that imposes no limits on the number of CpGs included in the model and effectively handles correlated variables. Therefore, the Elastic Net approach is a reasonable choice for the development of epigenetic clocks.

Instead of using DNA methylation ratios of CpGs as explanatory variables, recent proposals have suggested using variables transformed through principal component analysis (PCA) to estimate more reliable clocks (Higgins-Chen et al., 2022). PCA is a method that identifies orthogonal basis vectors, called principal components, from a data matrix, capturing more information with fewer dimensions. The transformation helps reduce technical noise introduced to explanatory variables during sample preparation and measurement. Higgins-Chen et al. (2022) re-estimated several epigenetic clocks: Horvath, Hannum, Levine (PhenoAge), Telomere length (DNAmTL), and GrimAge, based on PCA-transformed DNA methylation ratios, and reported that the prediction performances were comparable to or better than those of the original clocks (Horvath, 2013; Horvath and Raj, 2018; Hannum et al., 2013; Levine et al., 2018; Lu et al., 2019b,a; Higgins-Chen et al., 2022). These epigenetic clocks developed using PCA-transformed explanatory variables are referred to as PC clocks in this study.

Although many distinct epigenetic clocks have been developed, most have not been tailored to specific racial populations. Epigenetic status can be influenced by genetic or environmental factors in a particular population, potentially introducing biases in existing clocks when applied to them. To develop a biological age model that performs better for a specific population, it is necessary to collect DNA methylation data from that population (Hicken et al., 2023). Large sample sizes are required due to the high dimensionality of the data; for example, the PC clocks in Higgins-Chen et al. (2022) were trained using data from thousands of individuals. However, collecting such large sample sizes is challenging due to the high costs and time requirements. To address this issue, we considered using a transfer learning approach (Yang et al., 2020). Transfer learning enables the development of models from the data with a relatively small sample size by using information from related data or models from other populations. Therefore, in this study, we proposed a new transfer learning method that uses the parameter information of a model trained with the Elastic Net.

This study also focused on developing epigenetic clocks for the Japanese population, particularly PC clocks to obtain more reliable models. Previously, some epigenetic clocks were developed with Japanese data consisted of 421 registrants, which are designed to predict the chronological age (Komaki et al., 2023). Our study expands beyond the first-generation clock to include the development of a second-generation clock, Levine’s PhenoAge, for the Japanese population (Levine et al., 2018). PhenoAge is a prominent second-generation clock designed to predict the adjusted age for biomarkers related to age-associated mortality.

The remainder of this paper is organized as follows: Section 2 describes the proposed transfer learning method. Section 3 explains the development processes of epigenetic clocks for the Japanese population. Section 4 presents a comparison of the performance of the original clocks, clocks trained using conventional methods, and those trained using our proposed transfer learning approach. Section 5 discusses the methodology and results. Finally, Section 6 concludes the study.

## 2 Transfer Learning via Regularization

Suppose we have a dataset consisting of *n* observations, where *y* = (*y*_1_, …, *y*_*n*_)^⊤^ ∈ ℝ^*n*^ is the response variable and *X* = (*X*_1_, …, *X*_*n*_)^⊤^ ∈ ℝ^*n*×*p*^ is the matrix of explanatory variables. *X*_*i*_ ∈ ℝ^*p*^ is the vector of explanatory variables for observation *i*. Let *β* = (*β*_0_, …, *β*_*p−*1_)^⊤^ ∈ ℝ^*p*^ represent the regression coefficient vector.

Transfer learning involves a suite of techniques that utilize knowledge gained from addressing one or more problems to solve another related problem. Various approaches exist based on what information is transferred and how it is done under some assumptions. In this study, we assumed that while we can use the parameters of the regression model, we cannot use the actual data from the source domain, and we aimed to develop a regression model in the target domain using data that shares the same features as the source domain. We specifically focused on parameter transfer via regularization. We explored the use of parameter information from models estimated through sparse regularization methods in the source domain to estimate models for the related problem in the target domain. In the context of the epigenetic clock, the source domain involved a multiracial population, and the target domain was the Japanese population. Data from the source domain may not be readily available due to the need to protect subjects’ privacy.

### 2.1 Transfer Lasso

The Transfer Lasso was recently proposed by Takada and Fujisawa (2020) to facilitate the transfer of parameter information from a model trained using the least absolute shrinkage and selection operator (Lasso) (Tibshirani, 1996). The estimator of the Transfer Lasso is defined as:

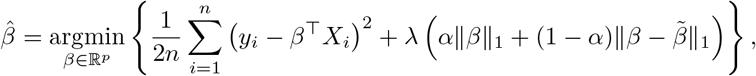

where 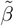 represents the initial estimate obtained from the source domain. The tuning parameters *λ ∈* [0, +*∞*) and *α ∈* [0, 1] control the intensity of the regularization and the balance between the two *ℓ*_1_-norm components, respectively. While the first component of the regularization term is the same as that in the traditional Lasso, the second component represents the difference between the parameters of the source and target, promoting sparsity in the changes in the target estimates from the source estimates, as well as the estimates themselves. Given that the Elastic Net, rather than the Lasso, is typically used for estimating epigenetic clocks, it is necessary to adapt the Transfer Lasso to the Elastic Net for our purposes (Zou and Hastie, 2005).

### 2.2 Transfer Elastic Net

We propose a Transfer Elastic Net, which straightforwardly extends the regularization terms of the Transfer Lasso to the transfer learning of Elastic Net. The proposed estimator is defined as:

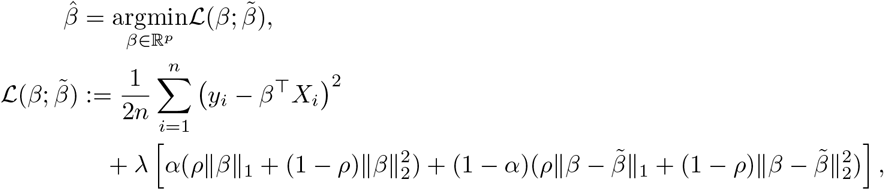

where 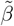 is the initial estimate, and λ ∈ [0, +∞), *α* ∈ [0, 1], and *ρ* ∈ [0, 1] are the tuning parameters. The additional tuning parameter, *ρ* controls the balance between the *ℓ*_1_ and *ℓ*_2_ norm components. This method effectively uses the parameter information of a model trained using the Elastic Net in the source domain to inform the learning process in the target domain.

We developed an estimation algorithm for the Transfer Elastic Net based on the coordinate descent.

By partially differentiating 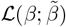 with respect to *β*_*j*_, we obtain:

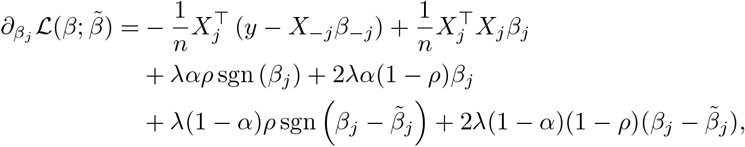

where the explanatory variable matrix without the *j*th vector *X*_*j*_ is denoted by *X*_−*j*_ ∈ ℝ^*n*×(*p*−1)^ and the parameter vector *β* without the *j*th component *β*_*j*_ is denoted by *β*_−*j*_ ∈ ℝ^*p*−1^. Solving the equation 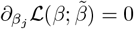 for *β*_*j*_ yields the following updating rule:

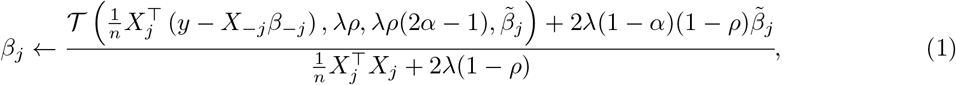

where 𝒯 is the soft-threshold function:

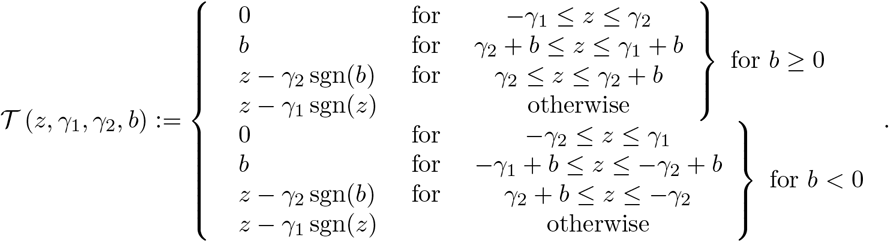

This soft-threshold function is identical to that used in Transfer Lasso. The update rule (1) for *j* = 0, …, *p −* 1 is applied iteratively until convergence is achieved, as detailed in Algorithm 1.

#### Algorithm 1

Estimation algorithm for Transfer Elastic Net

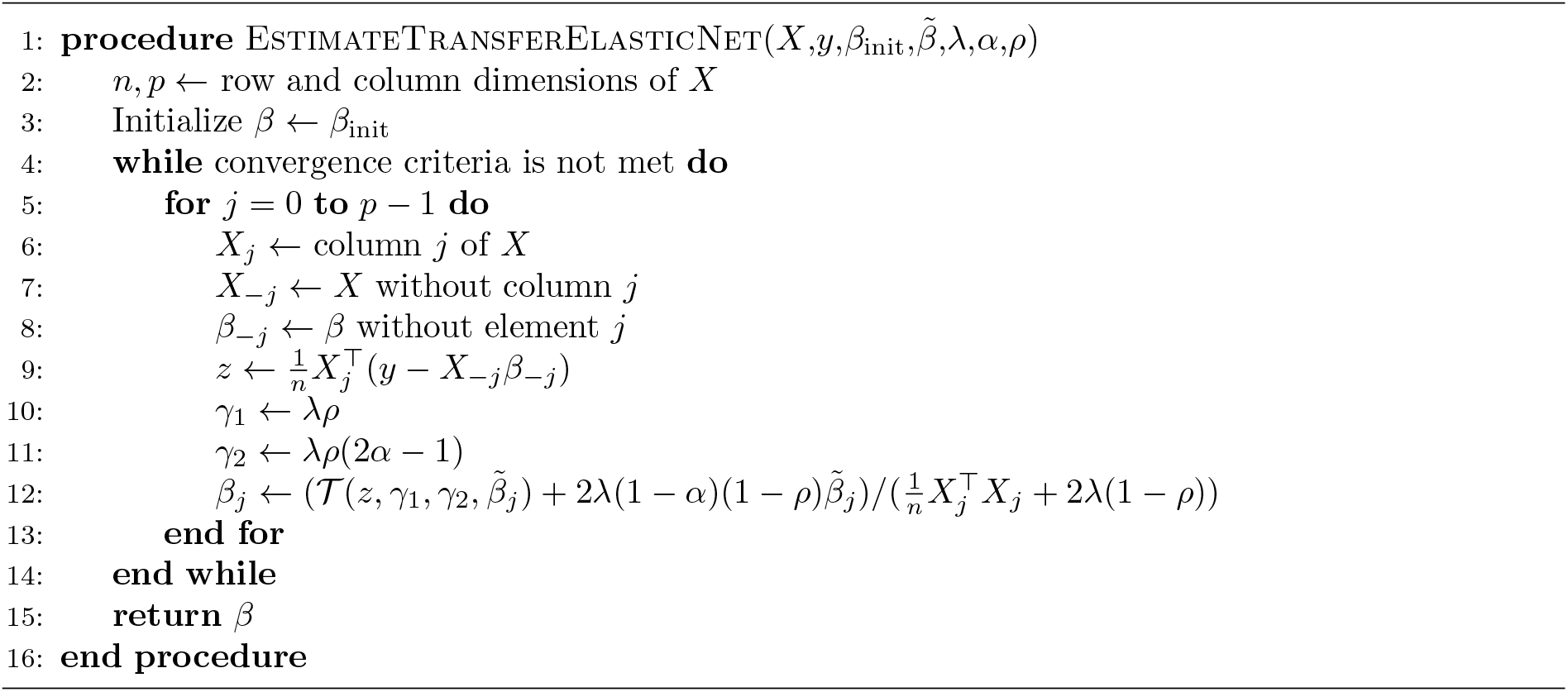

## 3 Development of PC-clocks for the Japanese Population

We developed Horvath’s, Hannum’s, and Levine’s (PhenoAge) clock types for the Japanese population using principal component-transformed DNA methylation data: PC-Horvath, PC-Hannum, and PC-PhenoAge, respectively, (Horvath, 2013; Hannum et al., 2013; Levine et al., 2018). This section outlines the data collection and processing, as well as the model training and evaluation procedures.

### 3.1 Data Collection

DNA methylation and blood biomarkers were measured from the blood samples of the 194 subjects included in our study. The study population comprised 76 healthy males and 118 females, aged 23–84 years, who did not require regular medication or treatment.

Genomic DNA was extracted from the blood samples using the Promega Maxwell® RSC Genomic DNA Kit. The obtained genomic DNA was treated with bisulphite using the Zymo Research EZ DNA Methylation Kit, following the standard protocol provided by Illumina, Inc. (hereafter referred to as “Illumina”). The treated DNA was amplified, fragmented, and hybridized to the Illumina Infinium Methylation EPIC BeadChip, adhering to Illumina’s specified procedures (https://support.illumina.com/array/array_kits/infinium-methylationepic-beadchip-kit/documentation.html). The arrays were imaged using the Illumina iScan System set to the Illumina’s recommended scanning parameters.

Then, the nine blood biomarkers were quantified: Albumin (g/L), Creatinine (umol/L), Serum Glucose (mmol/L), C-reactive protein (mg/dL), Lymphocyte percent (%), Mean cell volume (fL), Red cell distribution width (%), Alkaline phosphatase (U/L), and White blood cell count (1000 cells/uL). One female subject with insufficiently qualified results was excluded from the study. Summary statistics of the measured characteristics of the participants are presented in the Results section.

### 3.2 Data Processing

While the response variable for PC-Horvath and PC-Hannum was chronological age, for PC-PhenoAge it was phenotypic age, which accounts for the variability in risk and physiological conditions among people of the same chronological age (Levine et al., 2018). Phenotypic age was calculated for each subject using the same method as Levine et al. (2018), incorporating chronological age and nine blood biomarkers. For the *j*th individual, represented by the subscript, phenotypic age (PhenotypicAge) was computed as follows:

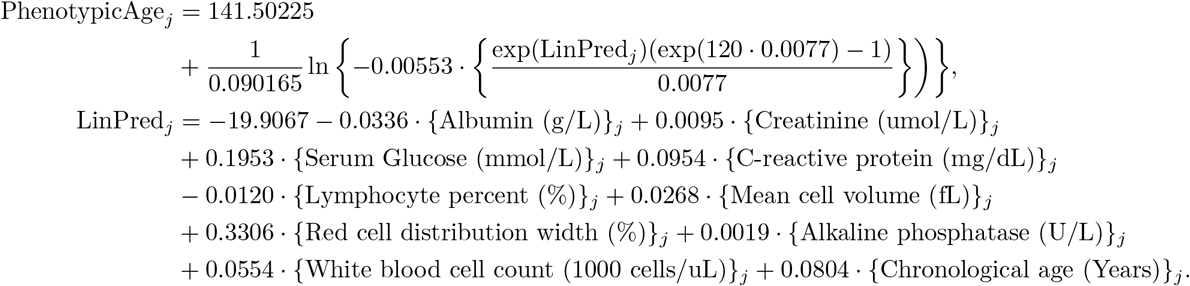

This calculation adjusts age for factors associated with aging-related mortality in NHANES III data, which are determined by the Cox proportional hazards model with *ℓ*_1_ regularization. Details are provided in the supplementary materials of Levine et al. (2018), and Lee and Wang (2003).

Raw IDAT data from the DNA methylation array were converted to DNA methylation ratios (beta values) for each probe using the ChAMP R package (version 2.24.0). Probe selection was based on stringent criteria: high quantitative validity (P value *<* 0.01 and bead counts *>* 5%), association with CpG sites distal to SNPs (Single Nucleotide Polymorphisms), unique mapping (not mapped to multiple locations), and exclusion from the X and Y chromosomes. The beta values for the selected probes were normalized using the BMIQ method (Teschendorff et al., 2013), implemented in the ChAMP R package. Additionally, batch effects due to variations in processing times were corrected using the ComBat function (Johnson et al., 2007), also implemented in the ChAMP R package.

We focused on principal component-based clocks to develop more reliable models, which use transformed values through the principal component analysis instead of raw methylation ratios for explanatory variables (Higgins-Chen et al., 2022). The data processing followed the workflow implemented in the PC-Clocks software (https://github.com/MorganLevineLab/PC-Clocks). Due to the absence of 6024 necessary CpGs in our data for principal component transformation using the PC-Clocks software, we imputed these missing values using the GSE40279 dataset (Hannum et al., 2013). We then centered and transformed the values using the trained principal component parameters specific to each clock prepared in the software. For the construction of the models, we used the first 4,282 principal components for PC-Horvath, 657 for PC-Hannum, and 4505 for PC-PhenoAge, as specified in the software and documented in the previous study (Higgins-Chen et al., 2022).

### 3.3 Training Methods and Processes

We used the conventional Elastic Net (ENet) and the proposed Transfer Elastic Net (TENet) approaches to train the clocks. The values of the tuning parameters were determined by 5-fold cross-validation. The candidate values for the tuning parameters of TENet were λ ∈ {0.0625, 0.125, 0.25, 0.5, 1}, *ρ* ∈ {0, 0.25, 0.5, 0.75, 1}, and *α* ∈ {0, 0.25, 0.5, 0.75, 1}. For ENet, the candidate values for *λ* and *ρ* were the same as for TENet; however, *α* was fixed at 1. The coefficients from the original clocks were used as the initial estimates, 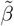 for TENet.

Of the 193 samples, 50 were randomly selected for the evaluation set, while the remaining 143 were used for the training set. We used the training set to determine the values of the tuning parameters and estimate the coefficients. Subsequently, we evaluated the performances of the trained models using the evaluation set.

We used Python 3.8.3 to perform the estimation process. The estimation algorithms for ENet and TENet were implemented using Numpy 1.23.3. The cross-validation procedure was conducted using scikit-learn 1.3.1 (Pedregosa et al., 2011).

### 3.4 Evaluation and Comparison

We compared the performances of the three models in Table 1 for PC-Horvath, PC-Hannum, and PC-PhenoAge. The evaluation metrics included root mean squared error (RMSE), mean absolute error (MAE), and Pearson correlation coefficient (PCC). Each estimated model was applied to the evaluation set, and the metrics were calculated. The relationships between predicted chronological or phenotypic age and the corresponding true values were presented through scatter plots. The number of nonzero coefficients for each combination of models was counted, and the cosine similarity between each pair of model parameters was calculated.

**Table 1:**
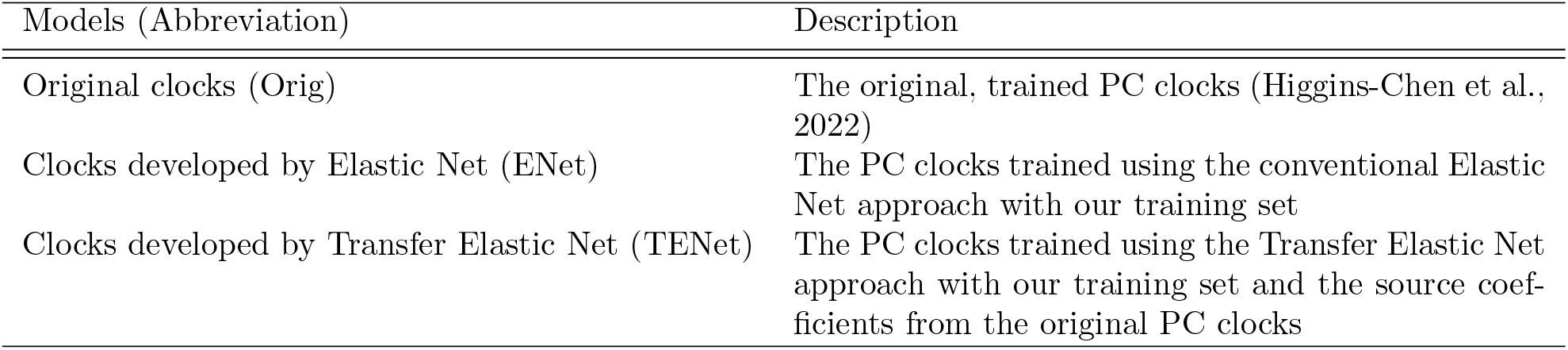
The compared models.

| Models (Abbreviation) | Description |
| --- | --- |
| Original clocks (Orig) | The original, trained PC clocks (Higgins-Chen et al., 2022) |
| Clocks developed by Elastic Net (ENet) | The PC clocks trained using the conventional Elastic Net approach with our training set |
| Clocks developed by Transfer Elastic Net (TENet) | The PC clocks trained using the Transfer Elastic Net approach with our training set and the source coefficients from the original PC clocks |

The evaluation metrics were calculated using scikit-learn 1.3.1, and plots were created using matplotlib 3.2.2.

## 4 Results

### 4.1 Characteristics of the Subjects

Summary statistics for the characteristics of the collected samples are presented in Table 2 separately for the training and evaluation sets. The training set comprised subjects aged 25–84 years in chronological age, whereas the evaluation set included subjects aged 23–82 years. Although the proportion of females was slightly higher in the evaluation set, few statistical differences were observed between the two datasets.

**Table 2:** Characteristics of the training and evaluation sets.

| Characteristic | Training set (143 samples) |  |  |  | Evaluation set (50 samples) |  |  |  |
| --- | --- | --- | --- | --- | --- | --- | --- | --- |
|  | Mean | SD | Min | Max | Mean | SD | Min | Max |
| Chronological Age (Year) | 54.83 | 12.99 | 25.81 | 84.66 | 51.91 | 14.72 | 23.61 | 82.76 |
| Phenotypic Age | 50.61 | 14.53 | 18.39 | 88.84 | 47.25 | 18.76 | 9.50 | 95.41 |
| Sex (Male:0, Female:1) | 0.58 | - | - | - | 0.68 | - | - | - |
| Albumin (g/L) | 42.37 | 3.43 | 35.62 | 50.00 | 42.90 | 3.70 | 37.94 | 56.00 |
| Creatinine (umol/L) | 70.68 | 18.36 | 35.36 | 144.98 | 66.53 | 17.53 | 41.55 | 124.64 |
| Serum Glucose (mmol/L) | 5.62 | 1.48 | 3.66 | 14.32 | 5.82 | 2.27 | 3.33 | 16.32 |
| C-reactive protein (mg/dL) | 0.08 | 0.16 | 0.03 | 1.33 | 0.10 | 0.17 | 0.03 | 1.19 |
| Lymphocyte percent (%) | 32.14 | 7.48 | 11.80 | 59.70 | 30.12 | 8.63 | 11.20 | 49.30 |
| Mean cell volume (fL) | 94.01 | 4.75 | 71.00 | 106.00 | 94.48 | 3.86 | 86.00 | 102.00 |
| Red cell distribution width (%) | 13.03 | 1.07 | 11.60 | 20.20 | 12.78 | 0.64 | 11.60 | 14.20 |
| Alkaline phosphatase (U/L) | 63.73 | 17.22 | 36.00 | 115.00 | 60.64 | 14.76 | 27.00 | 89.00 |
| White blood cell count (1000 cells/uL) | 5.40 | 1.29 | 2.10 | 9.40 | 5.41 | 1.69 | 3.00 | 12.60 |

### 4.2 Comparison Results

The evaluation results for the clocks are presented in Table 3. All clocks developed using our proposed TENet exhibited the smallest prediction errors in terms of RMSE and MAE and achieved the highest PCC values (PC-Horvath: RMSE of 3.91, MAE of 2.93, and PCC of 0.968; PC-Hannum: RMSE of 3.56, MAE of 2.86, and PCC of 0.971; PC-PhenoAge: RMSE of 7.58, MAE of 5.55, and PCC of 0.933). ENet’s performance ranked second except for the PCC in PC-PhenoAge (PC-Horvath: RMSE of 5.32, MAE of 4.16, and PCC of 0.954; PC-Hannum: RMSE of 3.96, MAE of 3.13, and PCC of 0.963; PC-PhenoAge: RMSE of 9.07, MAE of 6.82, and PCC of 0.888), while the Orig clocks typically performed the worst (PC-Horvath: RMSE of 6.60, MAE of 5.19, and PCC of 0.942; PC-Hannum: RMSE of 10.43, MAE of 9.21, and PCC of 0.957; PC-PhenoAge: RMSE of 9.19, MAE of 7.26, and PCC of 0.921).

**Table 3:**
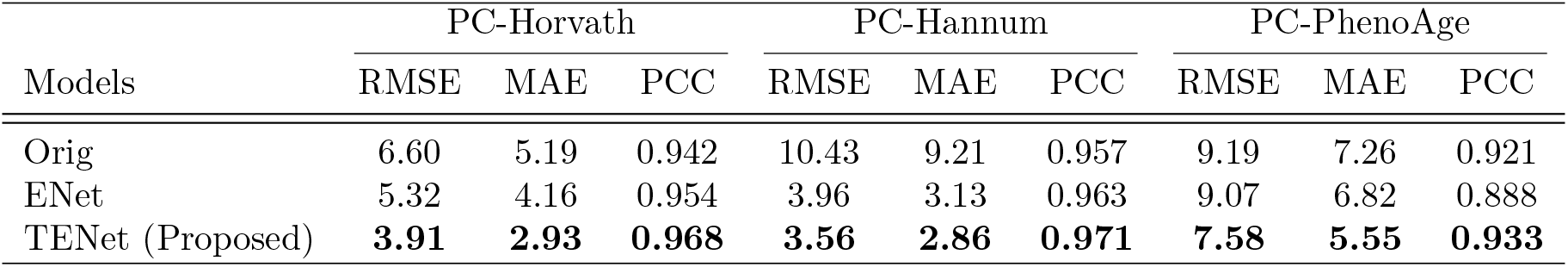
Prediction performance of the original and developed models.

| Models | PC-Horvath |  |  | PC-Hannum |  |  | PC-PhenoAge |  |  |
| --- | --- | --- | --- | --- | --- | --- | --- | --- | --- |
|  | RMSE | MAE | PCC | RMSE | MAE | PCC | RMSE | MAE | PCC |
| Orig | 6.60 | 5.19 | 0.942 | 10.43 | 9.21 | 0.957 | 9.19 | 7.26 | 0.921 |
| ENet | 5.32 | 4.16 | 0.954 | 3.96 | 3.13 | 0.963 | 9.07 | 6.82 | 0.888 |
| TENet (Proposed) | **3.91** | **2.93** | **0.968** | **3.56** | **2.86** | **0.971** | **7.58** | **5.55** | **0.933** |

The predicted chronological or phenotypic ages versus the true values in the evaluation set are plotted in Figure 1. For all PC clocks, Orig tended to overestimate the age of relatively young subjects. From the results of ENet, we observed that this bias was reduced when training with Japanese data. TENet further reduced both the bias and the variance around the true values.

**Figure 1.**
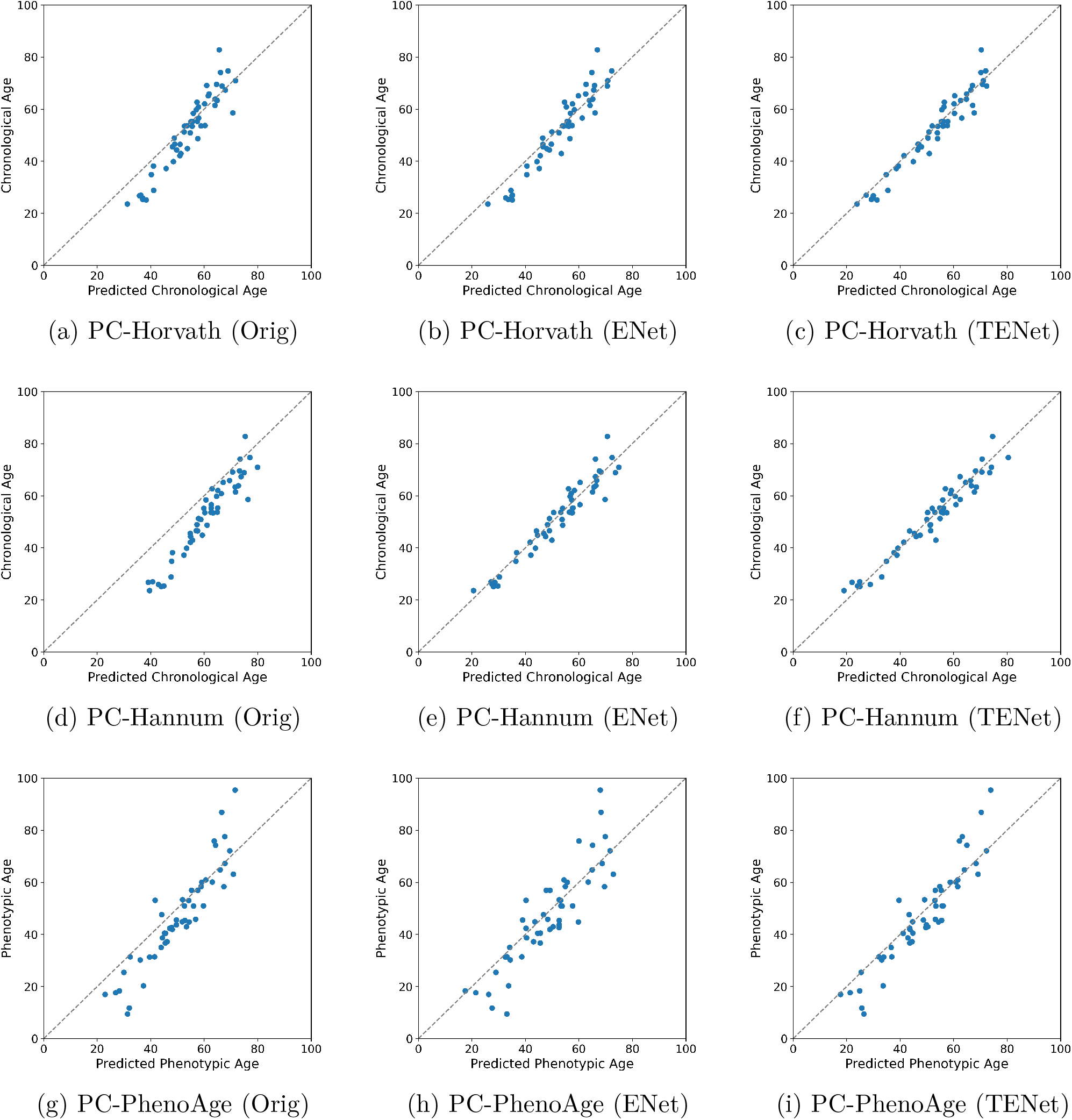
True vs. Predicted values of age in the evaluation set

The number of nonzero elements, the number of common nonzero elements for combinations, and the cosine similarity of the estimated coefficients (excluding the intercept) were calculated and are shown in Table 4. For PC-Horvath, the counts of nonzero elements for the Orig and TENet were both 120. For ENet, it was 4282, indicating that all the variables were selected in the estimated model. The combination of the Orig and TENet showed 120 overlapping nonzero elements and a higher cosine similarity than the combinations of the Orig and ENet, and TENet and ENet (0.828 for Orig&TENet, 0.589 for Orig&ENet, and 0.500 for TENet&ENet). For PC-Hannum and PC-PhenoAge, similar values were observed concerning the number of nonzero elements for the original and TENet (PC-Hannum: 389 for Orig, 390 for TENet, with 389 in common; PC-PhenoAge: 651 for Orig, 652 for TENet, with 651 in common), and the similarity of the coefficients (PC-Hannum: 0.795 for Orig&TENet, 0.683 for Orig&ENet, and 0.403 for TENet&ENet; PC-PhenoAge: 0.960 for Orig&TENet, 0.464 for Orig&ENet, and 0.366 for TENet&ENet). However, the number of nonzero elements for ENet’s coefficients was relatively small (PC-Hannum: 44; PC-PhenoAge: 20).

**Table 4:** Number of non-zero coefficients and similarity of coefficients from the trained models.

| PC-clocks | Models | non-Zeros | Common non-zeros | Cosine similarity |
| --- | --- | --- | --- | --- |
| PC-Horvath | Orig | 120 | - | - |
|  | TENet | 120 | - | - |
|  | ENet | 4282 | - | - |
|  | Orig & TENet | - | 120 | 0.828 |
|  | Orig & ENet | - | 120 | 0.589 |
|  | TENet & ENet | - | 120 | 0.500 |
|  | Orig & TENet & ENet | - | 120 | - |
| PC-Hannum | Orig | 389 | - | - |
|  | TENet | 390 | - | - |
|  | ENet | 44 | - | - |
|  | Orig & TENet | - | 389 | 0.795 |
|  | Orig & ENet | - | 38 | 0.683 |
|  | TENet & ENet | - | 38 | 0.403 |
|  | Orig & TENet & ENet | - | 38 | - |
| PC-PhenoAge | Orig | 651 | - | - |
|  | TENet | 652 | - | - |
|  | ENet | 20 | - | - |
|  | Orig & TENet | - | 651 | 0.960 |
|  | Orig & ENet | - | 19 | 0.464 |
|  | TENet & ENet | - | 20 | 0.366 |
|  | Orig & TENet & ENet | - | 19 | - |

## 5 Discussion

From the results, PC clocks for the Japanese population developed via the proposed transfer learning approach consistently performed better than the original clocks and those developed using the conventional estimation method. For all the clocks we developed, the parameters estimated via the Transfer Elastic Net were sparse and more similar to the parameters in the original models than those estimated with the standard Elastic Net. These results are consistent with the expected behavior of the Transfer Elastic Net, thereby supporting the efficacy of the proposed method.

Our results also showed that the prediction bias in younger Japanese groups observed in the original clocks was corrected when models were trained using Japanese data. However, due to the small sample size of the training dataset, the prediction variance of the standard Elastic Net approach was higher than that of the transfer learning approach. Furthermore, in the estimation using the conventional Elastic Net method, while all candidate variables were selected for PC-Horvath, only a very small number of variables were selected for PC-Hannum and PC-PhenoAge, indicating instability in the estimation processes. Our proposed approach, the Transfer Elastic Net, addresses these issues by estimating parameters to be close to zero or to the parameters of the source models, using the Japanese population dataset.

Due to differences in DNA methylation measurement methods, the epigenetic clocks for the Japanese population in Komaki et al. (2023) cannot be directly applied to our data. Thus, we cannot directly compare our clocks with theirs. However, through the proposed transfer learning approach, we successfully developed epigenetic clocks that outperform the original PC clocks in Higgins-Chen et al. (2022), using about half the sample size in Komaki et al. (2023), and achieved comparable Pearson’s correlation coefficients in the evaluation set (about 0.94 to 0.97 in their results). Moreover, PC-PhenoAge for the Japanese population is the first second-generation clock for the population to our knowledge. Therefore, our results remain significant.

Although our focus was on PC clocks in this study, it is also feasible to perform transfer learning using DNA methylation data without transformation. One potential advantage of using the raw methylation ratio of CpGs is the inclusion of the DNA methylation ratio of CpGs not measured in the source domain as additional candidate explanatory variables. Then, it may be possible to explore genetic factors associated with aging based on information about the selected CpGs. However, the estimation may not be stable, due to the high dimensionality of the explanatory variables.

There are many other areas of bioinformatics research, in addition to epigenetics, where collecting a large sample size is challenging due to the high costs or rarity of the samples. When using sparse regularization approaches for the development of prediction or discrimination models, such as models predicting phenotypes from gene expression or models discriminating outcomes from metabolome, even if the sample size available in the target domain is small, our proposed transfer learning approach may allow us to develop better-performing models if a related model in the source domain is available. As the Transfer Elastic Net is a general method for parameter transfer of Elastic Net, its application is not limited to the bioinformatics field.

In this study, we assumed that data in the source domain were not available; however, the parameters of the regression models were available. Therefore, we considered a parameter-transfer approach. Other transfer learning approaches for Elastic Net may be applicable if data in the source domain are available (Li et al., 2015; Handl et al., 2019).

The Elastic Net has some advantages over the Lasso, including the grouping effect (Zou and Hastie, 2005; Zhou, 2013). The grouping effect states that the coefficients of highly correlated explanatory variables have small differences, resulting in these variables being included or excluded from the estimated model simultaneously. Further investigation is required to determine whether Transfer Elastic Net exhibits similar properties. In addition, there are many variations in sparse regularization methods. For example, some approaches use non-convex regularization terms to reduce the shrinkage toward zero of coefficients remaining in the estimated model, such as bridge penalty, smoothly clipped absolute deviation (SCAD), and minimax concave penalty (MCP) (Frank and Friedman, 1993; Fan and Li, 2001; Zhang, 2010). Developing parameter transfer methods for these non-convex regularization methods is also a future work.

## 6 Conclusion

In conclusion, we proposed Transfer Elastic Net as a parameter transfer approach for Elastic Net, and we developed principal component-based epigenetic clocks for the Japanese population using the proposed method. Our results demonstrate that these epigenetic clocks perform better than the original clocks or those trained using conventional Elastic Net, despite using approximately half the sample size of the prior study. Our proposed transfer learning approach can be used to construct epigenetic clocks for other specific populations and may be further applied to other bioinformatics or various fields to develop prediction models with relatively small sample sizes for the dimensions of the variables.

## Acknowledgement

We extend our deepest gratitude to Mr. Tomoyuki Matsuno and Mr. Asuka Kubo of Rhelixa Inc., who facilitated the acquisition of the critical dataset for this study through diligent coordination with Y’s Science Clinic. We are profoundly thankful to Dr. Sawako Hibino, whose efforts in collecting Japanese blood samples formed the foundation of our data. Additionally, we wish to express our appreciation to Dr. Genki Yoshikawa, who processed the raw data from the DNA methylation array to generate the methylation rate data essential for conducting the development of our clocks. The contributions of these individuals were indispensable to the achievements of this study. We also thank Editage (www.editage.jp) for English language editing.

## Disclosure statement

This work was funded by Rhelixa Inc. YT served as a technical advisor in statistical science from April 2021 to March 2024. RN is the founder and the chief executive officer.

This study was approved by the Ethics Committee of the Asia-Oceania Anti-Aging Promoting Association [AOAAPA22-001].

## Author’s contributions

Conceptualization: YT, RN; Data collection: RN; Statistical Methodology: YT; Data Processing and Analysis: YT, RN; Discussion about the Results: YT, RN; Writing – original draft: YT; Writing – review & editing: YT, RN.

## Data Availability

The sets of estimated coefficients of the PC clocks developed in this study will be accessible in the public repository on GitHub (https://github.com/t-yui/TransferENet-EpigeneticClock) upon acceptance of this manuscript. The Python script for the Transfer Elastic Net, including the implementation of test cases, is available from the same GitHub repository. Individual-level data will be made available upon reasonable request.

## References

Alfonso, G. and Gonzalez, J. R. (2020). Bayesian neural networks for the optimisation of biological clocks in humans. bioRxiv.

Fan, J. and Li, R. (2001). Variable selection via nonconcave penalized likelihood and its oracle properties. Journal of the American Statistical Association, 96(456):1348–1360.

Frank, L. E. and Friedman, J. H. (1993). A statistical view of some chemometrics regression tools. Technometrics, 35(2):109–135.

Handl, L., Jalali, A., Scherer, M., Eggeling, R., and Pfeifer, N. (2019). Weighted elastic net for unsupervised domain adaptation with application to age prediction from DNA methylation data. Bioinformatics, 35(14):i154–i163.

Hannum, G., Guinney, J., Zhao, L., Zhang, L., Hughes, G., Sadda, S., Klotzle, B., Bibikova, M., Fan, J.-B., Gao, Y., et al. (2013). Genome-wide methylation profiles reveal quantitative views of human aging rates. Molecular Cell, 49(2):359–367.

Hicken, M. T., Dou, J., Kershaw, K. N., Liu, Y., Hajat, A., and Bakulski, K. M. (2023). Racial and ethnic residential segregation and monocyte DNA methylation age acceleration. JAMA Network Open, 6(11):e2344722–e2344722.

Higgins-Chen, A. T., Thrush, K. L., Wang, Y., Minteer, C. J., Kuo, P.-L., Wang, M., Niimi, P., Sturm, G., Lin, J., Moore, A. Z., et al. (2022). A computational solution for bolstering reliability of epigenetic clocks: Implications for clinical trials and longitudinal tracking. Nature Aging, 2(7):644–661.

Horvath, S. (2013). DNA methylation age of human tissues and cell types. Genome Biology, 14:1–20.

Horvath, S. and Raj, K. (2018). DNA methylation-based biomarkers and the epigenetic clock theory of ageing. Nature Reviews Genetics, 19(6):371–384.

Johnson, W. E., Li, C., and Rabinovic, A. (2007). Adjusting batch effects in microarray expression data using empirical bayes methods. Biostatistics, 8(1):118–127.

Komaki, S., Nagata, M., Arai, E., Otomo, R., Ono, K., Abe, Y., Ohmomo, H., Umekage, S., Shinozaki, N. O., Hachiya, T., et al. (2023). Epigenetic profile of japanese supercentenarians: a cross-sectional study. The Lancet Healthy Longevity, 4(2):e83–e90.

Lee, E. T. and Wang, J. (2003). Statistical methods for survival data analysis, volume 476. John Wiley & Sons.

Levine, M. E., Lu, A. T., Quach, A., Chen, B. H., Assimes, T. L., Bandinelli, S., Hou, L., Baccarelli, A. A., Stewart, J. D., Li, Y., et al. (2018). An epigenetic biomarker of aging for lifespan and healthspan. Aging (Albany NY), 10(4):573.

Li, Y., Vinzamuri, B., and Reddy, C. (2015). Constrained elastic net based knowledge transfer for healthcare information exchange. Data Mining and Knowledge Discovery, 29:1094–1112.

Lu, A. T., Binder, A. M., Zhang, J., Yan, Q., Reiner, A. P., Cox, S. R., Corley, J., Harris, S. E., Kuo, P.-L., Moore, A. Z., et al. (2022). DNA methylation grimage version 2. Aging (Albany NY), 14(23):9484.

Lu, A. T., Quach, A., Wilson, J. G., Reiner, A. P., Aviv, A., Raj, K., Hou, L., Baccarelli, A. A., Li, Y., Stewart, J. D., et al. (2019a). DNA methylation grimage strongly predicts lifespan and healthspan. Aging (Albany NY), 11(2):303.

Lu, A. T., Seeboth, A., Tsai, P.-C., Sun, D., Quach, A., Reiner, A. P., Kooperberg, C., Ferrucci, L., Hou, L., Baccarelli, A. A., et al. (2019b). DNA methylation-based estimator of telomere length. Aging (Albany NY), 11(16):5895.

Pedregosa, F., Varoquaux, G., Gramfort, A., Michel, V., Thirion, B., Grisel, O., Blondel, M., Prettenhofer, P., Weiss, R., Dubourg, V., Vanderplas, J., Passos, A., Cournapeau, D., Brucher, M., Perrot, M., and Duchesnay, E. (2011). Scikit-learn: Machine learning in Python. Journal of Machine Learning Research, 12:2825–2830.

Takada, M. and Fujisawa, H. (2020). Transfer learning via ℓ_1_ regularization. Advances in Neural Information Processing Systems, 33:14266–14277.

Teschendorff, A. E., Marabita, F., Lechner, M., Bartlett, T., Tegner, J., Gomez-Cabrero, D., and Beck, S. (2013). A beta-mixture quantile normalization method for correcting probe design bias in illumina infinium 450 k DNA methylation data. Bioinformatics, 29(2):189–196.

Tibshirani, R. (1996). Regression shrinkage and selection via the lasso. Journal of the Royal Statistical Society Series B: Statistical Methodology, 58(1):267–288.

Yang, Q., Zhang, Y., Dai, W., and Pan, S. J. (2020). Transfer Learning. Cambridge University Press.

Zhang, C.-H. (2010). Nearly unbiased variable selection under minimax concave penalty. The Annals of Statistics, 38(2):894 – 942.

Zhang, Q., Vallerga, C. L., Walker, R. M., Lin, T., Henders, A. K., Montgomery, G. W., He, J., Fan, D., Fowdar, J., Kennedy, M., et al. (2019). Improved precision of epigenetic clock estimates across tissues and its implication for biological ageing. Genome Medicine, 11(1):1–11.

Zhou, D.-X. (2013). On grouping effect of elastic net. Statistics & Probability Letters, 83(9):2108–2112.

Zou, H. and Hastie, T. (2005). Regularization and variable selection via the elastic net. Journal of the Royal Statistical Society Series B: Statistical Methodology, 67(2):301–320.

